# CD82 expression marks the endothelium to hematopoietic transition at the onset of blood specification in human

**DOI:** 10.1101/2023.02.27.530202

**Authors:** Sara Menegatti, Bethany Potts, Roberto Paredes, Eva Garcia-Alegria, Syed Murtuza Baker, Valerie Kouskoff

**Affiliations:** Developmental Hematopoiesis Group, Faculty of Biology, Medicine and Health, the University of Manchester, Manchester M13 9PT, UK; CytoSeek Ltd, Unit Dx, Albert Road, Bristol, BS2 0XJ, UK; Division of Informatics, Imaging & Data Sciences, Faculty of Biology, Medicine and Health, the University of Manchester, Manchester M13 9PT, UK

## Abstract

**SUMMARY:** During embryonic development, all blood progenitors are initially generated from endothelial cells that acquire a hemogenic potential. Blood progenitors emerge through an endothelial-to-hematopoietic transition regulated by the transcription factor RUNX1. To date, we still know very little about the molecular characteristics of hemogenic endothelium and the molecular changes underlying the transition from endothelium to hematopoiesis. Here, we analysed at the single cell level a human embryonic stem cell-derived endothelial population containing hemogenic potential. RUNX1-expressing endothelial cells, which harbour enriched hemogenic potential, show very little molecular differences to their endothelial counterpart suggesting priming toward hemogenic potential rather than commitment. Additionally, we identify CD82 as a marker of the endothelium-to-hematopoietic transition. CD82 expression is rapidly upregulated in newly specified blood progenitors then rapidly downregulated as further differentiation occurs. Together our data suggest that endothelial cells are first primed toward hematopoietic fate, then rapidly undergo the transition from endothelium to blood.

## INTRODUCTION

During embryonic development, the hematopoietic system is established in sequential waves occurring at specific times and in specific anatomic locations, a feature highly conserved across evolution (Dzierzak and Bigas, 2018; Palis, 2016). In vertebrates, the first wave of blood specification is initiated in the yolk sac and gives rise to primitive erythroid, macrophage and megakaryocyte progenitors (Palis et al., 1999). A second wave of specification, still within the yolk sac, generates definitive erythroid and myeloid progenitors, shortly followed by the emergence of lymphoid progenitors (McGrath et al., 2015). The third and final wave of blood specification occurs within the major arteries of the embryo (de Bruijn et al., 2000). This last wave gives rise to embryonic multipotent progenitors (Patel et al., 2022) and hematopoietic stem cells (HSCs) that colonize the foetal liver (Medvinsky and Dzierzak, 1996). These embryonic progenitors and HSCs, homing to the bone marrow before birth, are the founders of the adult hematopoietic system and maintain the production of all blood cells throughout life (Costa et al., 2012).

In vertebrates, during embryonic development, blood stem and progenitor cells are initially generated from endothelial cells that acquire a hemogenic potential and are termed hemogenic endothelium (HE) (Lacaud and Kouskoff, 2017). Blood progenitors emerge from HE through an endothelial-to-hematopoietic transition regulated to a large extent by the evolutionary conserved transcription factor RUNX1 (Chen et al., 2009; Gao et al., 2018; Lancrin et al., 2009). Mouse embryos deficient for RUNX1 expression die by mid-gestation with a complete absence of all blood cells, except for primitive erythrocytes (Okuda et al., 1996; Wang et al., 1996). Similarly, both mouse and human RUNX1 deficient embryonic stem cells (ESCs) do not produce blood cells upon differentiation, with the exception of primitive erythrocytes (Bruveris et al., 2020; Lacaud et al., 2002). While these studies have established the essential role of RUNX1 for blood cell emergence, the molecular changes underlying the transition from endothelium to hematopoiesis still remains poorly understood. Additionally, we still know very little about the molecular characteristics of HE as the hemogenic potential of endothelial cells can only be defined retrospectively in further cultured to reveal or not hematopoietic potential.

Here, we analysed at the single cell level a human ESC-derived endothelial population containing hemogenic potential. RUNX1-expressing endothelial cells, which harbour enriched hemogenic potential, show very little molecular differences to their endothelial counterpart suggesting priming toward hemogenic potential rather than commitment. Additionally, we identify CD82 as a marker of endothelium-to-hematopoietic transition. CD82, a member of the tetraspanin 4 superfamily, is rapidly upregulated in newly specified blood progenitors and downregulated as soon as the further differentiation occurs.

## RESULTS

### Transcriptomic homogeneity of the CD31^+^CD144^+^ progenitor population

We previously identified the CD31^+^CD144^+^ cell population generated by day 6 of hESC differentiation in serum-free culture (Figure S1A) as the cell population most enriched for hemogenic endothelium giving rise to hematopoiesis (Garcia-Alegria et al., 2018). To further characterise this CD31^+^CD144^+^ cell population in an unbiased approach, single-cell RNA sequencing was performed using the 10X Genomics platform. The small CD43^+^ fraction within this CD31^+^CD144^+^ cell population was kept to provide an internal reference of committed hematopoietic cells within the dataset (Figure 1A). Following data quality check and filtering (Figure S1B-S1E), a total of 2,537 cells and 11,818 genes were identified for downstream analysis. Dimensionality reduction with t-distributed stochastic neighbour embedding (t-SNE) and clustering with dynamic-tree cut resulted in the identification of 12 clusters (Figure 1B). An immediate observation was the broad overlap between clusters and the absence of clearly defined populations, except for cluster 12. Uniform Manifold Approximation and Projection (UMAP) and Force-directed graph were used as additional dimensionality reduction approaches (Figure 1C and 1D). However, clusters separation was not improved, suggesting a relative homogeneity within the transcriptional landscape of the cells analysed. Accordingly, the identification of differentially regulated genes for each cluster versus all other clusters revealed a very limited number of genes upregulated with Log2 fold change greater than 2 in most clusters except for clusters 9, 11 and 12 (Table 1 and Figure S2). The expression of most endothelial genes such as *KDR*, *CDH5* and *SOX7* was uniformly distributed across most clusters (Figure 2A and S3A). In contrast, a few genes were markedly differentially expressed, among which the *CXCR4* arterial gene was predominantly expressed in cluster 2 and 6 and *ESM1* implicated in angiogenesis and neovascularisation (Vila-Gonzalez et al., 2019) was mostly expressed in cluster 2 (Figure 2B). However, no clear clustering of arterial or venous markers was observed (Figure S3B), with most cells expressing both types of genes. Despite applying cell cycle-effect removal filters, a subset of genes relating to cell cycle, such as *HIST1H1D* or *AURKB*, was more predominantly expressed in cluster 5 (Figure 2B-C and S3C). Based on their relative proximity on t-SNE plot, some of the clusters were grouped and further analysed. When all upregulated genes of cluster 1, 2 and 6 were intersected, 302 genes were found commonly upregulated which enriched for GO terms such as angiogenesis, cell adhesion and extracellular matrix organization (Figure 2D). On the other hand, cluster 3, 4 and 5 had only 55 commonly upregulated genes, mainly related to cell division (Figure 2E). Together, these data suggest that the CD31^+^CD144^+^ cell population is a fairly homogenous pool of early endothelial cells with no specification yet toward arterial or venous fate. Cells within this population seem to differ only slightly by their state of endothelial identity or proliferation.

**Table 1:** Top 10 differentially upregulated genes per cluster (Cluster x versus all remaining clusters) sorted by Log2 Fold Change. Genes that are upregulated with Log2 fold change (FC) greater than 2 are highlighted in red.

| Cluster 1 |  | Cluster 2 |  | Cluster 3 |  | Cluster 4 |  |
| --- | --- | --- | --- | --- | --- | --- | --- |
| Gene | Log<sub>2</sub>FC | Gene | Log<sub>2</sub>FC | Gene | Log<sub>2</sub>FC | Gene | Log<sub>2</sub>FC |
| S100A4 | 1.02 | ESM1 | 3.29 | APLN | 1.13 | CCND1 | 1.19 |
| CAV1 | 0.87 | HPGD | 2.39 | CCND1 | 0.89 | NRP2 | 1.14 |
| CD300LG | 0.81 | C10orf10 | 1.83 | TM4SF18 | 0.85 | ZAP70 | 1.11 |
| S100A3 | 0.76 | CXCR4 | 1.69 | FAM198B | 0.77 | KCNMB1 | 1.02 |
| RNASE1 | 0.74 | IGFBP3 | 1.67 | CXorf36 | 0.71 | GRAP2 | 0.90 |
| GJA4 | 0.68 | PRND | 1.52 | SOX17 | 0.70 | APLN | 0.90 |
| UNC5B | 0.64 | GJA1 | 1.44 | HOPX | 0.66 | TPBGL | 0.87 |
| NOTCH4 | 0.60 | A2M | 1.44 | ESAM | 0.61 | FZD10 | 0.86 |
| PRKCDBP | 0.59 | KCNN3 | 1.33 | THY1 | 0.55 | PIM3 | 0.85 |
| EFNB2 | 0.59 | UNC5B | 1.30 | RHOC | 0.53 | APLNR | 0.85 |

| Cluster 5 |  | Cluster 6 |  | Cluster 7 |  | Cluster 8 |  |
| --- | --- | --- | --- | --- | --- | --- | --- |
| Gene | Log<sub>2</sub>FC | Gene | Log<sub>2</sub>FC | Gene | Log<sub>2</sub>FC | Gene | Log<sub>2</sub>FC |
| HIST1H4C | 2.55 | CARTPT | 4.53 | IGFBP5 | 2.99 | COL3A1 | 3.31 |
| HIST1H1D | 2.38 | SOX6 | 1.61 | ANXA1 | 1.66 | CRHBP | 2.47 |
| HIST1H1B | 2.28 | LPL | 1.57 | LYPD6B | 1.64 | BST2 | 2.03 |
| RRM2 | 1.97 | EPAS1 | 1.54 | BST2 | 1.63 | BEX1 | 1.92 |
| RAD51AP1 | 1.68 | KCNK17 | 1.48 | CRYBA4 | 1.54 | CTSV | 1.86 |
| CLSPN | 1.68 | PRSS23 | 1.43 | HOXB8 | 1.49 | GATA5 | 1.82 |
| ATAD2 | 1.62 | A2M | 1.42 | CACNA2D1 | 1.45 | CPED1 | 1.76 |
| HIST1H1E | 1.53 | CXCR4 | 1.41 | PLCG2 | 1.45 | HAND2 | 1.75 |
| TK1 | 1.50 | KCNN3 | 1.39 | HOXB.AS3 | 1.44 | SAMSN1 | 1.74 |
| HIST2H2AC | 1.49 | PLCG2 | 1.32 | ADAMTS6 | 1.44 | BTK | 1.74 |

| Cluster 9 |  | Cluster 10 |  | Cluster 11 |  | Cluster 12 |  |
| --- | --- | --- | --- | --- | --- | --- | --- |
| Gene | Log<sub>2</sub>FC | Gene | Log<sub>2</sub>FC | Gene | Log<sub>2</sub>FC | Gene | Log<sub>2</sub>FC |
| GYPB | 3.98 | TRGC1 | 2.12 | HAND1 | 5.99 | APOA2 | 7.95 |
| MYC | 3.89 | SCG5 | 2.11 | CER1 | 5.38 | APOA1 | 7.50 |
| GYPE | 3.63 | SELE | 1.87 | ACTC1 | 5.30 | RP11.148B6. | 5.95 |
| KLF1 | 3.50 | FN1 | 1.81 | KRT19 | 4.87 | S100A14 | 5.58 |
| SPN | 3.17 | LHX1 | 1.69 | LUM | 4.28 | POU5F1 | 5.54 |
| GATA1 | 3.10 | SFRP2 | 1.68 | MYL7 | 4.13 | LINC01356 | 5.27 |
| KCNH2 | 2.96 | ID4 | 1.56 | LIX1 | 4.08 | EPCAM | 4.99 |
| PDE8B | 2.94 | RIPPLY3 | 1.54 | RP11.834C11 | 4.03 | APELA | 4.93 |
| GYPA | 2.90 | ETV2 | 1.54 | HAS2 | 3.82 | TTR | 4.63 |
| GATA2 | 2.81 | NEFH | 1.49 | PRRX1 | 3.75 | RSPO3 | 4.55 |

**Figure 1:**
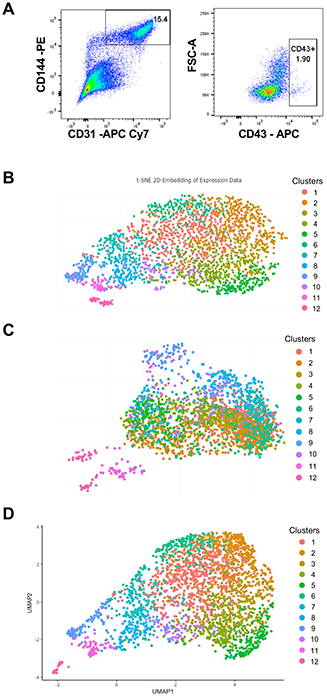
Characterisation of the CD31^+^CD144^+^ cell population from day 6 EBs. (A) FACS plot showing the sorted CD31^+^CD144^+^ cell population and the frequency of CD43^+^ cells contained within the CD31^+^CD144^+^ cell population used for single-cell RNA sequencing. (B) t-SNE plot of single cell RNA-seq data representing 12 clusters identified by dynamic-tree cut clustering. (C) Plot generated using UMAP for dimensionality reduction. (D) Plot generated using force-directed graph for dimensionality reduction.

**Figure 2:**
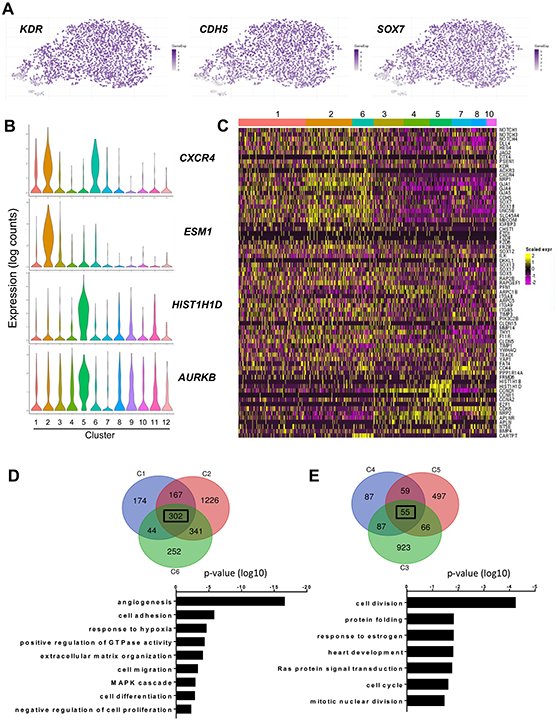
Low level of heterogeneity amongst endothelial cell clusters. (A) t-SNE plots showing the expression of *KDR*, *CDH5* and *SOX7* across all clusters. (B) Violin plots showing the differential expression of the arterial marker *CXCR4*, the angiogenic marker *ESM1* and the cell cycle-related genes *HIST1H1D* and *AURKB*. (C) Heatmap of differentially expressed genes within all endothelial clusters (1,2,3,4,5,6,7,8,10). The gene list includes arterial, venous, cell cycle-related genes, along with factors of Wnt, Notch and HIPPO pathways. (D) Venn diagram representing the number of genes upregulated in cluster 1, 2 and 6 with below the gene ontology analysis of all genes commonly upregulated in these three clusters. (E) Venn diagram representing the number of gene upregulated in cluster 3, 4 and 5 with below the gene ontology analysis of all genes commonly upregulated in these three clusters.

### Single cell RNA-seq identifies subsets of committed cells

While most clusters displayed strong endothelial characteristics, three small clusters showed commitment toward other lineages. The expression of genes related to the cardiomyocyte lineage, including *HAND1*, *ACTC1* or *MESP1*, was strongly enriched in cells from cluster 11 (Figure S4A). Gene ontology analysis confirmed enrichment in pathways related to cardiac muscle and heart development, although bone development-related terms were also enriched (Figure S4C). Cells in cluster 12 expressed epithelial-related markers, such as *EPCAM*, but also cardiac-related genes such as *TNNT1* and *APELA* (Figure S4B). Cluster 12 comprised two sub-clusters expressing each a few specific genes; the significance of these two sub-clusters is unclear (Figure S4D). The top gene ontology term for genes upregulated in cluster 12 cells was retinoid metabolic process which is known to play an important role in epicardial development (Wang and Moise, 2019) (Figure S4E). Together, these data suggest that cluster 12 contains an early population of epicardium progenitors as the expression of more mature epicardial genes such as *WT1* or *TBX18* was not observed (Niderla-Bielinska et al., 2019).

The expression of hematopoietic genes was mostly restricted to cells within cluster 9 (Figure 3A) with cells in the outer tip of this cluster enriched for *GATA1*, *SPN* or *SPI1* expression (Figure 3B). Gene set enrichment analysis (GSEA) of all genes upregulated in cluster 9 confirmed enrichment in hematopoietic cell lineage and cell cycle signature, associated with a decrease in the expression of adhesion molecules (Figure 3C). Gene ontology analysis confirmed enrichment in pathways related to hematopoiesis, erythrocyte and myeloid differentiation (Figure S5A). To identify possible trajectory and cell dynamics within this cluster, pseudotime was computed using *KDR* and *SPN* as anchor genes (Figure 3D). Pseudotime ordering revealed a single trajectory from cells expressing high levels of endothelial genes, including *KDR, CDH5* and *SOX7* to cells expressing genes indicative of hematopoietic commitment such as *GATA1, GYPA* and *SPN*. The expression of *GFI1* and *GFI1B*, two transcriptional repressors known to play a critical role in murine endothelial-to-hematopoietic transition (Lancrin et al., 2012; Thambyrajah et al., 2016), was observed in cells scattered throughout the pseudotime trajectory while *RUNX1* was expressed almost uniformly in all cells along this pseudotime. Unsupervised hierarchical clustering of selected endothelial and hematopoietic genes revealed a similar trend (Figure S5B). This pattern of expression is in line with the trajectory of cells undergoing an endothelial-to-hematopoietic transition in which the transcriptional landscape shifts from an endothelial-dominated program to a hematopoietic-committed program. Cluster 9 was thus classified as hematopoietic cluster, containing cells undergoing endothelial-to-hematopoietic transition and early committed hematopoietic cells.

**Figure 3:**
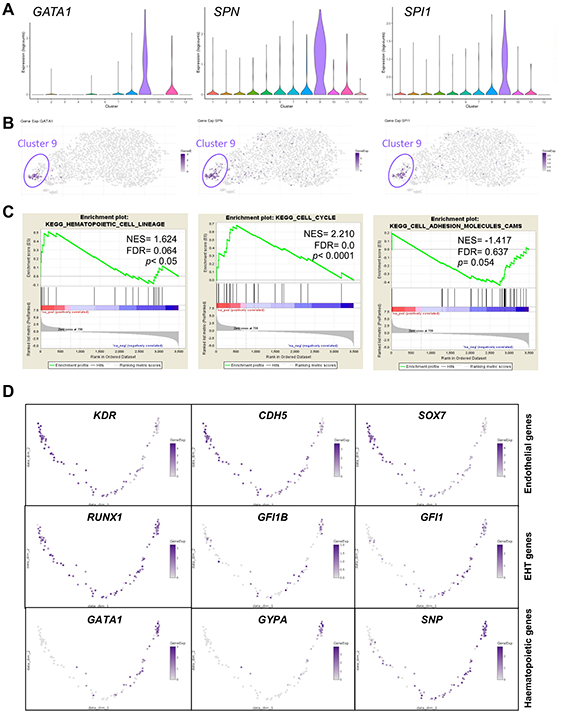
Characterisation of the hematopoietic cluster. (A) Violin plots showing the distribution of *GATA1*, *SPN* and *SPI1* expression in each cluster. (B) t-SNE plots showing the expression of *GATA1*, *SPN* and *SPI1* across all clusters. (C) Gene set enrichment analysis (GSEA) of cluster 9 differentially expressed genes showing positively and negatively enriched pathways. (D) Diffusion maps of all cells contained within cluster 9 ordered using *KDR* and *SPN* as left and right anchor genes, respectively. Pseudotime trajectories are shown for the expression of the indicated genes.

### RUNX1 expression marks a subset of endothelial progenitors

We previously showed that the expression of RUNX1 enriches for hemogenic potential in both mouse (Sroczynska et al., 2009) and human (Menegatti et al., 2021). Therefore, we next investigated the expression of *RUNX1* in the CD31^+^CD144^+^ scRNA-seq dataset. In addition to its broad expression in cluster 9, *RUNX1* was found expressed in cells scattered throughout all the endothelial clusters (Figure 4A), with slight increased frequencies in cluster 6 and 7 (Figure 4B). However, the expression of *RUNX1* did not define a specific cluster within the CD31^+^CD144^+^ endothelial population. To explore the identity of *RUNX1*-expressing endothelial cells, we next compared the transcriptomic landscape of *RUNX1*-positive and *RUNX1*-negative cells, excluding cells from cluster 9, 11 and 12 in this analysis (Figure 4C). Surprisingly, no striking difference was observed between the two cell populations; however, the expression of some arterial genes including *CXCR4*, *EFNB2* or *GJA4* was increased in *RUNX1*-positive cells while the expression of some venous genes, including *APLNR*, *NRP2* and *APLN*, was decreased (Figure 4C and S5C). However, when analysed by qPCR or flow cytometry, none of these genes showed statistically significant differences in expression between the CD31^+^CD144^+^ RUNX1^pos^ and RUNX1^neg^ fractions isolated from day 6 EBs generated with a *RUNX1b∷VENUS* reporter hESC line as previously described (Menegatti et al., 2021). Together, these data suggest that at this early stage of hemogenic endothelium specification, a small subset of CD31^+^CD144^+^ endothelial cells has upregulated RUNX1b expression but this has not yet led to significant changes in their transcriptional landscape.

**Figure 4:**
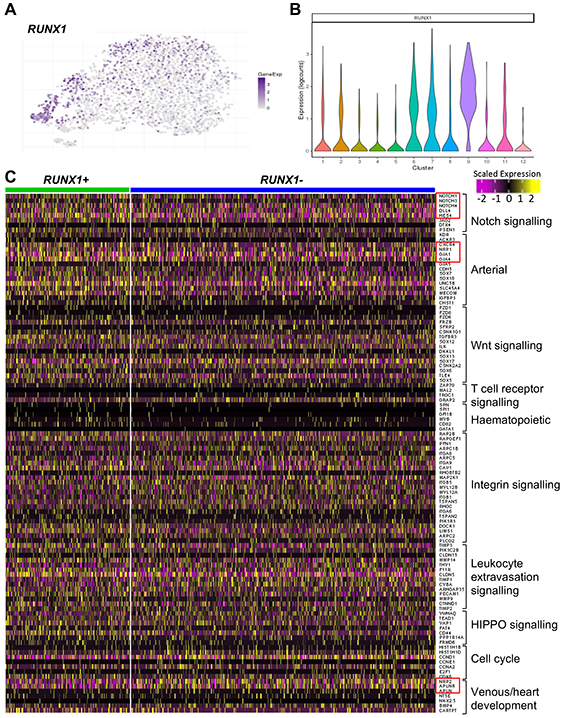
*RUNX1* expression is scattered throughout the endothelial clusters. (A) t-SNE plot showing *RUNX1* expression across all clusters. (B) Violin plot showing the distribution of *RUNX1* expression in each cluster. (C) Heatmap of the expression of genes involved in various pathways of interest as displayed on the right. Cells were divided in two clusters based on *RUNX1* expression. Cells with RUNX1 expression higher than zero are in the green cluster and cells with RUNX1 expression equal to zero are in the blue cluster. Cells included in this analysis belong to all clusters except for clusters 9, 11 and 12.

### CD82 expression marks the endothelium to hematopoietic transition

To further understand the molecular characteristic of the endothelium-to-hematopoietic transition, we next investigated the expression of genes differentially expressed in cells from cluster 9 relative to all other clusters, as these genes should mark the transition from endothelium to hematopoiesis. Most of the genes downregulated were indicative of endothelial identity loss (Supplemental table 1), previously shown to be controlled by the transcriptional repressors GFI1 and GFI1b in association with RUNX1 (Thambyrajah et al., 2016). We focused our attention on the genes most upregulated in cluster 9 (Figure 5A), marking the initiation of the hematopoietic program. On t-SNE plot representation, most of these genes were upregulated in a subset of cells located within the outer tip of cluster 9 (Figure S6). The expression of these genes marked commitment to the erythroid lineage, including glycophorin genes (*GYPA*, *GYPB*, *GYPE*) and transcription factors important for specification to erythroid (*KLF1*, *GATA1*, *NFE2*, *GFI1*) and myeloid lineages (*GATA2*, *SPI1*, *MYB*). Very few genes showed a pattern of expression throughout cluster 9 with the notable exception of CD82 (figure 5B), a member of the tetraspanin 4 superfamily. Of interest, CD82 is expressed on HSCs (Balise et al., 2020) and has recently been implicated in maintaining HSCs dormancy (Hur et al., 2016b) and preventing their mobilization (Saito-Reis et al., 2021).

**Figure 5:**
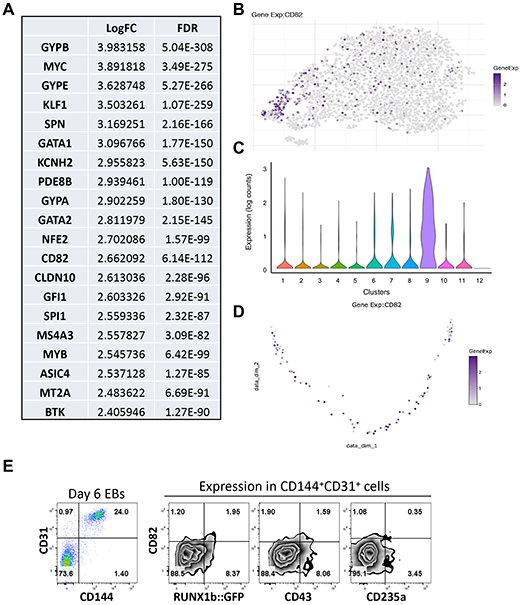
*CD82* expression is upregulated in cells from cluster 9. (A) List of the top 20 genes upregulated in cells from cluster 9 compared to all other clusters. (B) t-SNE plot showing *CD82* expression across all clusters. (C) Violin plot showing the distribution of *CD82* expression in each cluster. (D) Diffusion maps of all cells contained within cluster 9 ordered using *KDR* and *SPN* as left and right anchor genes, respectively. Pseudotime trajectory is shown for the expression of *CD82*. (E) Representative flow cytometry plots for the indicated cell surface markers in the CD31^+^CD144^+^ gated population at day 6 of EB differentiation of *RUNX1b∷VENUS* hESCs. Flow cytometry data are representative of three independent experiments.

Both t-SNE and Violin plots confirmed CD82 high expression in cluster 9 and low expression in all other clusters (Figure 5B and C). Pseudotime analysis of cluster 9 cells further showed CD82 upregulation as cells progressed from endothelium to hematopoiesis with a decreasing expression as cells became committed to erythroid (Figure 5D and 3D). To confirm the scRNA-seq data, CD82 expression was investigated by flow cytometry in the CD144^+^CD31^+^ population at day 6 of EB differentiation (Figure 5E). At this stage of differentiation, a small population of cells co-expressing low level of CD82 and RUNX1b was detected.

To determine the dynamic of CD82 expression upon hematopoietic specification, we analysed its expression alongside other markers over a 4-day culture of CD144^+^CD31^+^CD43^−^ isolated from day 6 EB (Figure 6A). Initially CD82 marked emerging cells positive for CD235a, CD43 and CD41a, but increasing frequencies of these blood committed cells progressively lost CD82 expression. By day 4 of the culture, more than half of CD43^+^ cells did not expressed CD82. In contrast, RUNX1 and CD82 remained co-expressed throughout the culture. This co-expression of the proteins was further confirmed by immuno fluorescence staining. Interestingly, RUNX1^+^ adherent cells mostly expressed CD82 intracellularly (Figure 6B) while free floating RUNX1^+^ cells showed a cell surface expression of CD82 (Figure 6C). Intrigued by the dynamic of CD82 expression, we next assessed the hematopoietic potential of cell populations expressing high, low or no CD82 (Figure 7A) These three populations were sorted from CD31^+^CD144^+^CD43^−^RUNX1∷VENUS^+^ cells isolated at day 6 of EB differentiation and culture for 4 days in hematopoietic inducing culture. We observed an increased hematopoietic potential that correlated with increasing CD82 expression level (Figure 7B, C). The CD82^−^ cell population mostly gave rise to small primitive erythrocyte colonies; the CD82^low^ cell population gave rise to fewer erythroid colonies harbouring a higher proliferation potential and a limit number of colonies with myeloid potential. The CD82^high^ cell population was the most enriched in high proliferative colonies, mostly of myeloid potential (Figure 7B, D).

**Figure 6:**
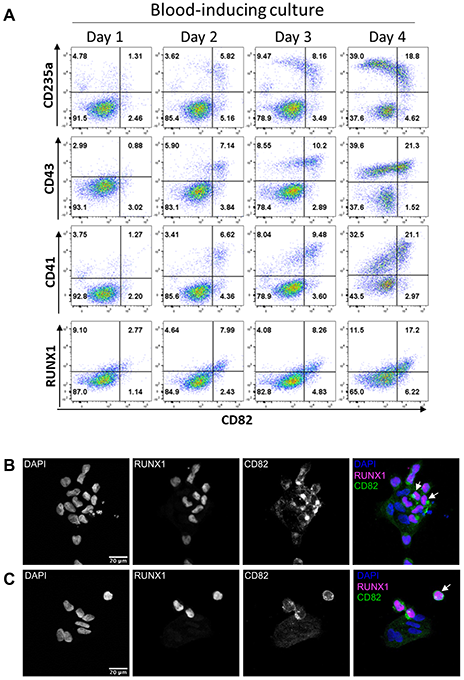
Dynamic of CD82 expression during blood cell emergence. (A) Representative flow cytometry plots for the indicated cell surface markers at day 1, 2, 3 and 4 of blood cell emergence from CD31^+^CD144^+^CD43^−^ cells isolated from day 6 of EB differentiation of *RUNX1b∷VENUS* hESCs and cultured in hematopoietic inducing condition. Flow cytometry data are representative of three independent experiments. (B) Representative photograph of immunofluorescence analysis for the indicated protein on CD31^+^CD144^+^CD43^−^ cells isolated from day 6 of EB differentiation of *RUNX1b∷VENUS* hESCs and cultured on Ibidi slides in hematopoietic inducing condition for two days. (C) Representative photograph of immunofluorescence analysis for the indicated protein on CD31^+^CD144^+^CD43^−^ cells isolated from day 6 of EB differentiation of *RUNX1b∷VENUS* hESCs and cultured on Ibidi slides in hematopoietic inducing condition for four days. All immunofluorescent staining data are representative of at least three independent experiments.

**Figure 7:**
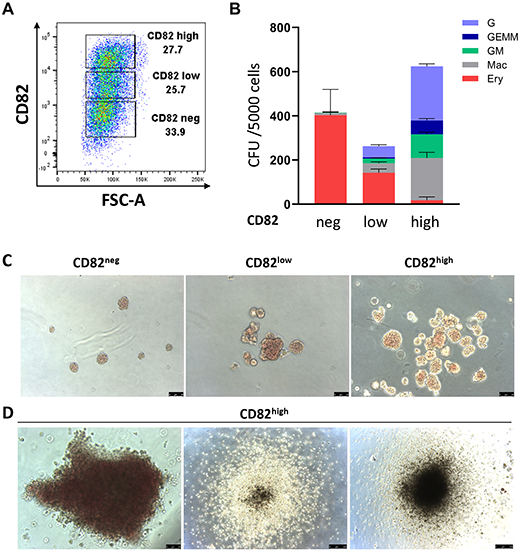
CD82 expression marks highly proliferative progenitors. (A) Representative flow cytometry plots showing the gating strategy for sorting CD82^neg^, CD82^low^ and CD82^high^ cell populations from CD31^+^CD144^+^CD43^−^RUNX1∷VENUS^+^ cells isolated from day 6 of EB differentiation of *RUNX1b∷VENUS* hESCs and cultured in hematopoietic inducing condition for 4 days. (B) CFU data for the indicated sorted populations, data are presented as mean +/− SD from three independent experiments performed in triplicate. G: granulocyte colonies; GEMM: granulocyte, erythrocyte, monocyte, megakaryocyte colonies; GM: granulocyte, monocyte colonies; Mac: macrophage colonies, Ery: erythrocyte colonies. (C) Representative photographs of erythrocyte colonies obtain in the CFU assays, pictures taken at day 7 of the CFU assay. (D) Representative photographs of high proliferative colonies from the CFU culture of the CD82^high^ population. Pictures taken at day 16 of the CFU assay. Scale bar: 50μM for all photographs in C and left photograph in panel D; scale bar: 250μM for centre and right photographs in panel D.

Together, these data revealed the transient expression pattern of CD82 at the onset of blood cell specification and uncover CD82 as a previously unrecognized marker of all hematopoietic progenitors in *in vitro* differentiating hESCs.

## DISCUSSION

All blood progenitors initially derive from endothelial cells with hemogenic properties. Understanding what makes an endothelium hemogenic is an area of intense investigation. Using scRNA-seq, we show here that the transcriptome of endothelial cells expressing RUNX1, which marks hemogenic endothelium (Bee et al., 2010; Menegatti et al., 2021; Sroczynska et al., 2009), has no overt distinctive molecular features when compared to the endothelial cells not expressing this transcription factor. However, endothelial cells that initiate the endothelium-to-hematopoietic transition up regulate CD82 expression. This cell surface marker remains expressed on the most immature blood progenitors but is downregulated as soon as further blood lineage differentiation occurs.

The transcriptomic analysis of the CD144^+^CD31^+^ cell population at the single cell level provides insightful information on the overall level of homogeneity of cells within this cell population. Most cells within this population have a strong endothelial identity and only differ by the expression of a few genes implicated in cell cycle or in specific endothelial function. It is not clear whether this represents a dynamic of gene expression that fluctuates randomly from cell to cell or whether subsets of endothelial cells are already specified or primed for specific functions. In the case of RUNX1-expressing endothelial cells, we observed increased expression of arterial markers and decreased expression of venous markers. However, when analysed at protein levels, there was no detectable expression or no differences between RUNX1 positive and negative cells, at least for the markers tested (CXCR4, DLL4, NRP1, and NRP2). This suggests that RUNX1-expressing endothelial cells might be primed toward an arterial fate and the activation of the Notch pathway. This is in line with recent findings from the Slukvin group (Park et al., 2018; Uenishi et al., 2018) but at odd with data published by Ditadi et al., suggesting that HE and arterial vascular endothelium represent distinct lineages (Ditadi et al., 2015).

The expression of CD82 upon transition from the endothelial to the hematopoietic program is an interesting finding, as CD82 is a well-known suppressor of cell motility and cancer metastasis (Tsai and Weissman, 2011; Vences-Catalan and Levy, 2022). Furthermore, CD82 is a critical regulator of hematopoietic progenitor migration and adhesion within the bone marrow (Balise et al., 2020; Hur et al., 2016b; Saito-Reis et al., 2018). The DARC molecule expressed by macrophages within the bone marrow niche has been shown to interact with CD82 at the cell surface of HSCs (Hur et al., 2016a; Hur et al., 2016b) to promote quiescent. On the other hand, several studies have demonstrated the importance of macrophages and pro-inflammatory signalling for blood cell emergence during embryonic development (Collins et al., 2021; Espin-Palazon et al., 2014; Frame et al., 2020; He et al., 2015; Li et al., 2014; Yang et al., 2022). It is tempting to speculate that in developing embryos, CD82 might contribute to the temporal maintenance of emerging blood progenitors in specific cellular niches in undifferentiated states.

### Limitations of the study

While we identified CD82 as a marker of endothelial to hematopoietic transition with an expression restricted to the most proliferative and multipotent blood progenitors, the functionality of CD82 remains elusive. CD82 knockout mice do not show any obvious defects or histopathologic abnormalities (Risinger et al., 2014). This lack of phenotype might be explained by compensatory mechanisms mediated by other members of the large family of Tetraspanin (Vences-Catalan and Levy, 2022) such as CD9 or CD151 that are also expressed by endothelial and blood cells (Balise et al., 2020; Zhang et al., 2009).

## Material and methods

### Single-cell RNA sequencing

For single-cell isolation, Man5 hESC cells were differentiated in EB culture for 6 days, dissociated and stained as previously described (Garcia-Alegria et al., 2021) and summarized below. Cell sorter FACS Aria Fusion was used to isolate live CD31^+^ CD144^+^ cells, which were then resuspended in PBS with 0.4% Bovine Serum Albumin (Sigma) at a density of 800 cells/μl. Cells were immediately loaded in the 10x-Genomics Chromium and libraries were prepared according to manufacturer’s guidelines, using Chromium Single Cell 3’ Solution (10X Genomics). Library sequencing was performed on Illumina NextSeq 500, with the aim of 100,000 reads per cell. Sequencing data were mapped onto GRCh38 human reference genome and quantified using Cell-Ranger Software (10x Genomics). For all downstream analyses listed below, R package was operated. Cells with read counts and number of expressed genes below 3 Median Absolute Deviation (MAD) were considered low quality and thus filtered out from the downstream analysis. Also, cells with percent of mitochondrial reads above 4 MAD were filtered out. We also removed the low-abundance genes from downstream analysis, by setting a 0.05 threshold for the number of cells in which each gene is expressed. Cell cycle phase classification was performed according to the prediction method described by Scialdone and colleagues (1) and the effect of this cell cycle was removed by considering it as a blocking factor during the identification of Highly Variable Genes (HVGs). Normalization of cell-specific biases was achieved using the deconvolution-based method (2) and log-transformation was applied (3), to avoid domination of downstream analysis by high-abundance genes with high variances. Normalized log-expression was then used to identify Highly Variable Genes (HVGs), which were thus applied for dimensionality reduction in t-distributed Stochastic Neighbour Embedding (t-SNE) (4), Uniform Manifold Approximation and Projection (UMAP) (5) and force-directed graph (6). For cell clustering, dynamicTreeCut R package was employed (7). Differentially expressed genes were analysed to identify marker genes for each of the appointed clusters. Cells were ordered also along pseudotime for trajectory prediction by inputting DE genes between different clusters identified by Monocle R package (8). Differentially expressed KDR and SPN genes were used as anchor points to direct pseudo temporal ordering, since these are known as the earliest and the latest marker genes expressed along endothelial to hematopoietic transition, respectively (9). Cells with RUNX1 expression higher than 0 were later isolated from the dataset for further cluster identification.

### ESCs maintenance and differentiation

Human ESCs were thawed and maintained on mitotically inactivated MEFs in Knock-Out DMEM media (Thermo Fisher Scientific) supplemented with 20% Knock-Out Serum Replacement (Thermo Fisher Scientific), 1% Minimum Essential Medium (MEM) Non-Essential Amino acid (Thermo Fisher Scientific), 2mM L-Gln, 25U/ml Pen/Strep, 0.1mM 2-mercaptoethanol and 8ng/ml human recombinant bFGF (PeproTech). Before differentiation, hESCs were feeder-depleted by culturing on Geltrex (Thermo Fisher Scientific) for 5 days in TeSR-E8 media (STEMCELL Technologies), supplemented with 25U/ml Pen/Strep. To generate EBs, hESCs were treated with EDTA and gently dissociated with EZPassage Tool (Thermo Fisher Scientific). Cell clumps were resuspended in StemPro-34 (Gibco) supplemented with 2mM L-Gln, 50U/ml Pen/Strep, 150ug/ml Transferrin, 50ug/ml Ascorbic Acid, 4.5 × 10 ^−4^ M MTG, Geltrex (1:200), 10uM ROCK inhibitor, 10ng/ml BMP4, plated in low-attachment dishes and incubated at 37°C 5% O_2_ 5% CO_2_. After 24 hours, 5ng/ml bFGF was added to each dish. At day 2, EBs were collected and resuspended in stemPro-34 supplemented with 2mM L-Gln, 50U/ml Pen/Strep, 150ug/ml Transferrin, 50ug/ml Ascorbic Acid, 4.5 × 10 ^−4^ M MTG, 10ng/ml BMP4, 5ng/ml bFGF, 0.9ng/ml Activin A. At day 4, EBs were collected and resuspended in stemPro-34 supplemented with 2mM L-Gln, 50U/ml Pen/Strep, 150ug/ml Transferrin, 50ug/ml Ascorbic Acid, 4.5 × 10 ^−4^ M MTG, 5ng/ml bFGF, 12ng/ml VEGF in StemPro-34). Further details are available in a Star Protocol (Garcia-Alegria et al., 2021).

### Hemogenic endothelium sort and hematopoietic inducing culture

After 6 days of culture, EBs were collected, disaggregated and stained with APC-eF780 conjugated anti-human CD31 (Thermo Fisher Scientific), PE-conjugated anti-human CD144 (BioLegend), PerCP-eF710-conjugated anti-human CD43 (Thermo Fisher Scientific) and Hoechst 33258 (Thermo Fisher Scientific). Live CD31^+^CD144^+^CD43^−^ cells were sorted and replated for in StemSpan (STEMCELL Technologies) supplemented with 5ng/ml VEGF, 5ng bFGF, 25ng/ml IGF1, 25ng/ml IGF2, 50ng/ml SCF, 50ng/ml TPO, 5ng/ml IL-11, 20ng/ml Flt3-L (all cytokines from PeproTech). Colony forming units (CFU) were performed using MethoCult™ SF H4636 (Stem Cell Technologies).

### Immunofluorescence analysis

Sorted CD144^+^CD31^+^CD43^−^ endothelial cells were grown on gelatin-coated glass slides (IBIDI) for 2 or 4 days in hematopoietic inducing conditions then fixed with 4% formaldehyde, blocked and permeabilized with 5% goat serum and 0.3% Triton-X100. Mouse monoclonal anti-CD82 antibody (Thermo Fisher cat no MA5-28570) was used at 1:100 and detected with an anti-mouse Alexa555 (Invitrogen) at 1:1,000. RUNX1 expression was either detected via VENUS expression from the RUNX1b∷VENUS allele or using the anti-RUNX antibody (ab92336, Abcam) at 1:1,000 detected with an anti-rabbit Alexa647 (Invitrogen) at 1:1,000. Both signals were previously shown to correlate accurately (Menegatti et al., 2021). Prolong Diamond antifade with DAPI (P36062, Invitrogen) was used as a mounting medium. Imaging acquisition was performed on a Leica SP8 inverted confocal microscope system using an HC PL APO CS2 40X/1.30 oil lens. 25 confocal planes (Z) were sequentially acquired for DAPI, VENUS and ALEXA555 emissions at 512×512, 16-bit pixels resolution and saved as .lif files.

## ACKNOWLEDGMENTS

The authors thank the staff at the Flow Cytometry, Bio-imaging and Genome Technology Core facilities of the University of Manchester for technical support. Research in the authors’ laboratory is supported by the Medical Research Council (MR/P000673/1), the Biotechnology and Biological Sciences Research Council (BB/R007209/1) and Cancer Research UK (C5759/A20971).

## AUTHOR CONTRIBUTIONS

Conceptualization: S.M. and V.K.; methodology: S.M. and V.K.; investigation: S.M., B.P., R.P. and E.G.A.; formal analysis: S.M. and S.M.B.; writing: S.M. and V.K.; supervision: E.G.A. and V.K.; Funding Acquisition, V.K.

