## supplemental figures for "CD82 expression marks the endothelium to hematopoietic transition at the onset of blood specification in human"

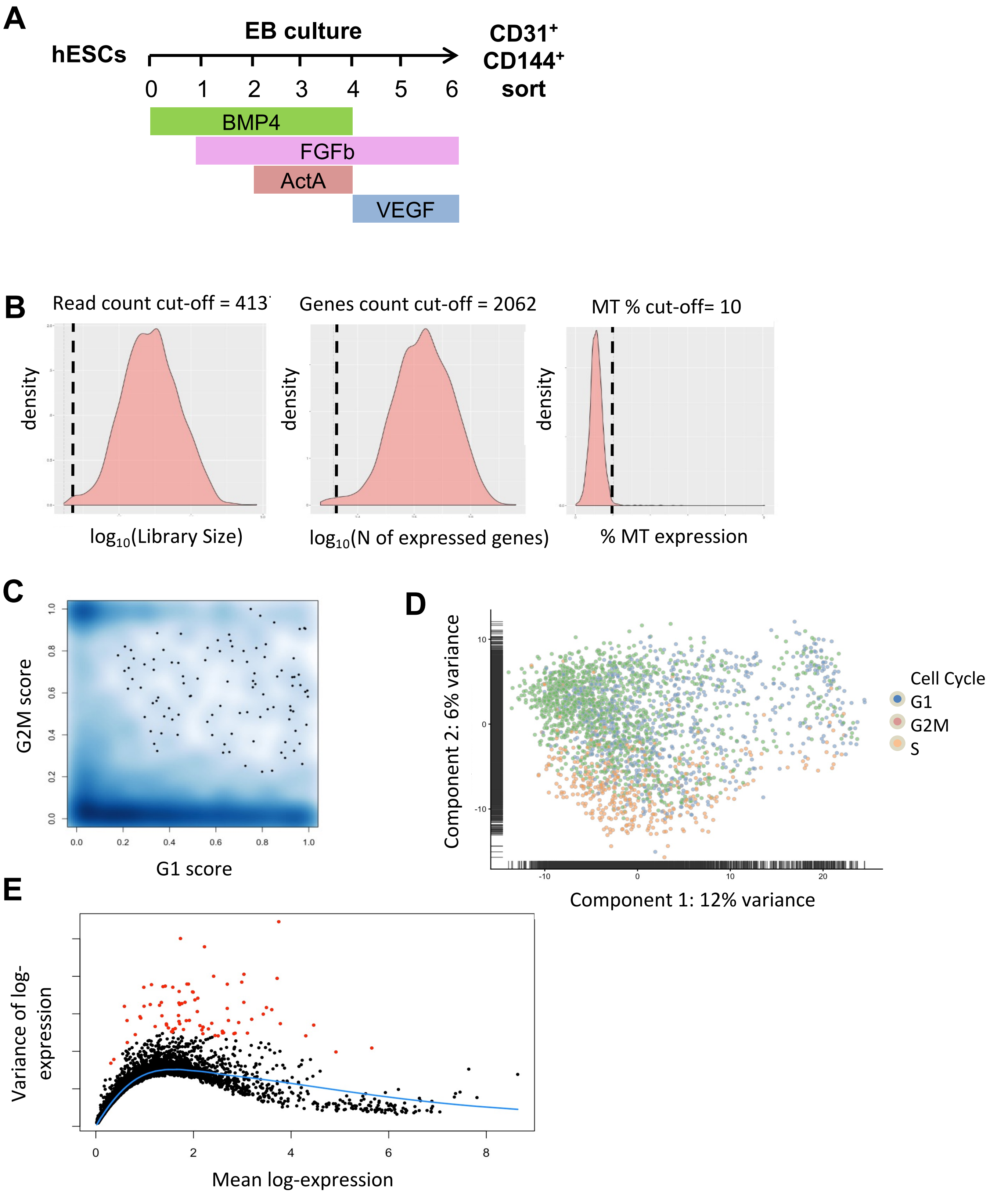
A
EB culture
hESCs
0 1 2 3 4 5 6
CD31+ CD144+ sort
BMP4
FGFb
ActA
VEGF
B
Read count cut-off = 413
Genes count cut-off = 2062
MT % cut-off= 10
density
log10(Library Size)
log10(N of expressed genes)
% MT expression
C
G2M score
G1 score
0.0 0.2 0.4 0.6 0.8 1.0
D
Component 2: 6% variance
Component 1: 12% variance
-10 0 10 20
Cell Cycle
G1
G2M
S
E
Variance of log-expression
Mean log-expression
0 2 4 6 8

Figure S1

**Figure S1. Differentiation scheme and scRNA-seq quality control.**

(A) Scheme of hESC differentiation toward endothelial and haematopoietic progenitors; hESCs are cultured as 3D embryoid bodies for 6 days, with the step-wise addition of growth factors. At day 0, EB media is supplemented with 10ng/ml BMP4. At day1, 5ng/ml FGFb is added to the culture. At day2, EB media is refreshed and supplemented with BMP4, FGFb and 0.9ng/ml Activin A. At day 4, medium is refreshed and supplemented with FGFb and 12ng/ml VEGF. (B) Normal distribution of cell density for library size, number of expressed genes and mitochondrial gene expression. Dotted lines define cut-offs used to filter low-quality cells. (C) Cell cycle analysis on generated dataset. Dark areas represent high cell density. (D) PCA on generated dataset showing cell cycle classification for each cell. (E) Identification of Highly Variable Genes (HVGs, red dots).

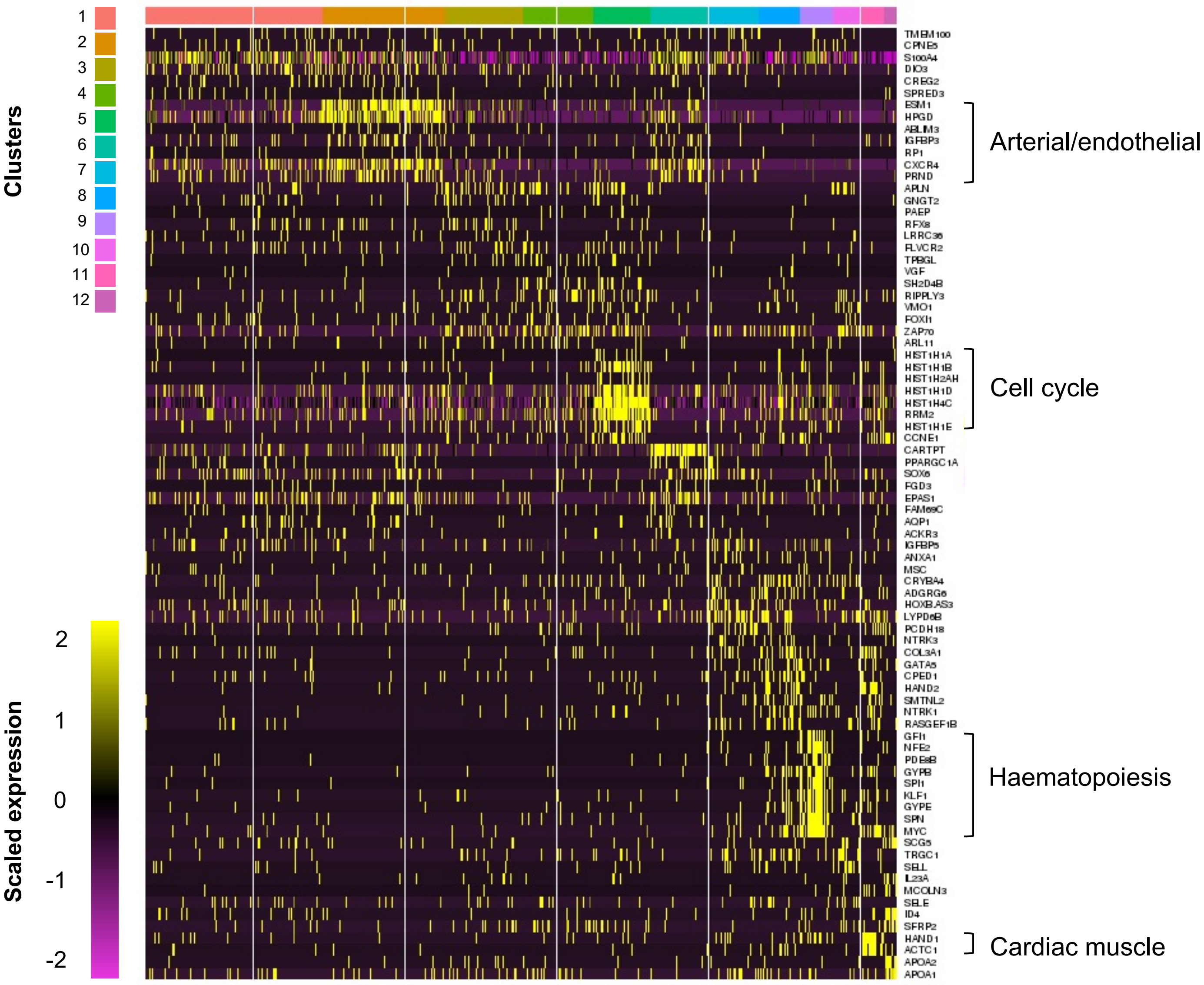


**Figure S2: Heatmap of the 10 most upregulated genes in each cluster.**
Grouping of arterial, cell cycle, haematopoiesis and cardiac muscle development are identifying some of the clusters included in the dataset.

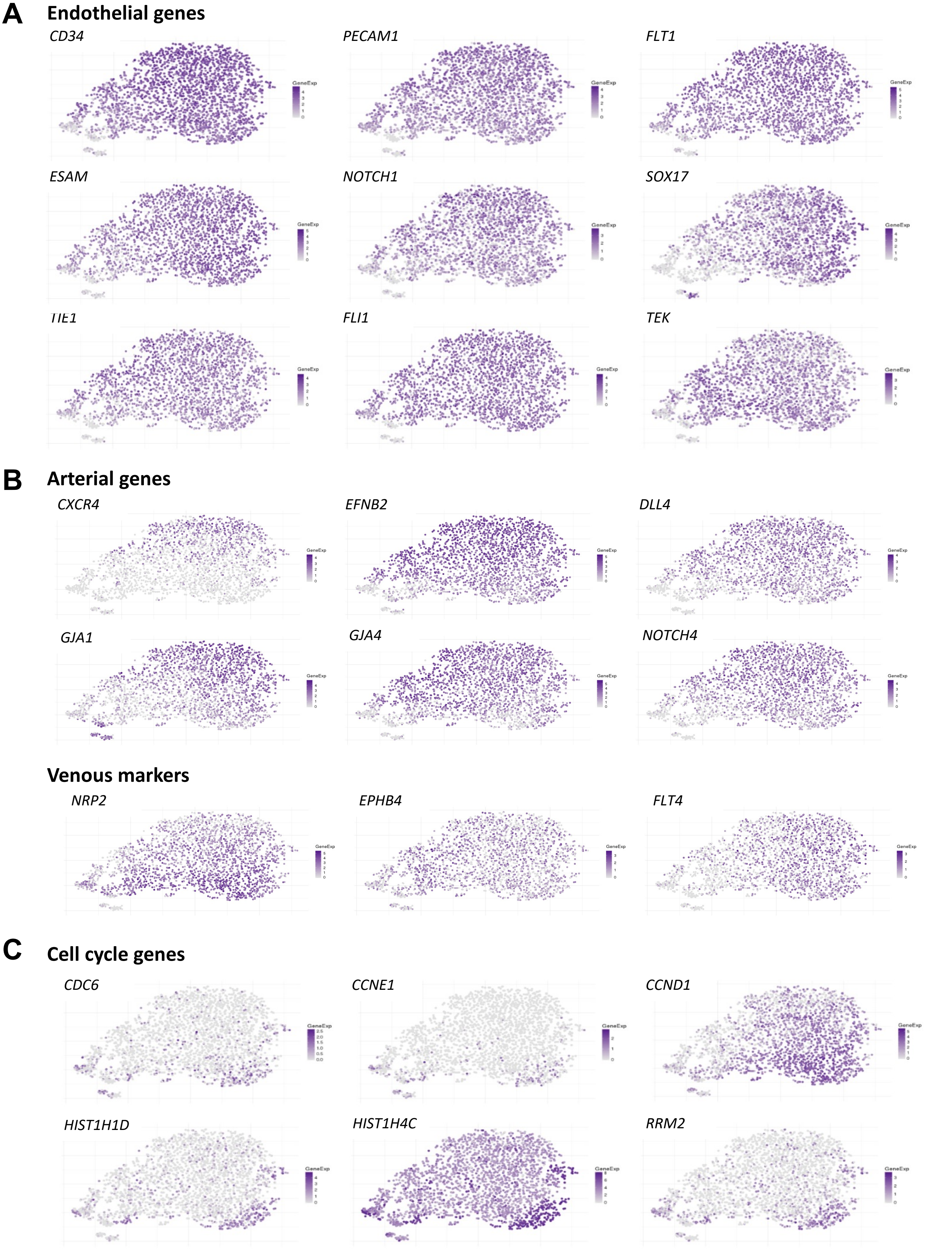
A
Endothelial genes
CD34
PECAM1
FLT1
ESAM
NOTCH1
SOX17
TIE1
FLI1
TEK
B
Arterial genes
CXCR4
EFNB2
DLL4
GJA1
GJA4
NOTCH4
Venous markers
NRP2
EPHB4
FLT4
C
Cell cycle genes
CDC6
CCNE1
CCND1
HIST1H1D
HIST1H4C
RRM2
GeneExp

Figure S3

**Figure S3. Gene expression across clusters.**

t-SNE plots showing the expression of endothelial genes (A), arterial and venous genes (B) and cell cycles genes (C) across all clusters.

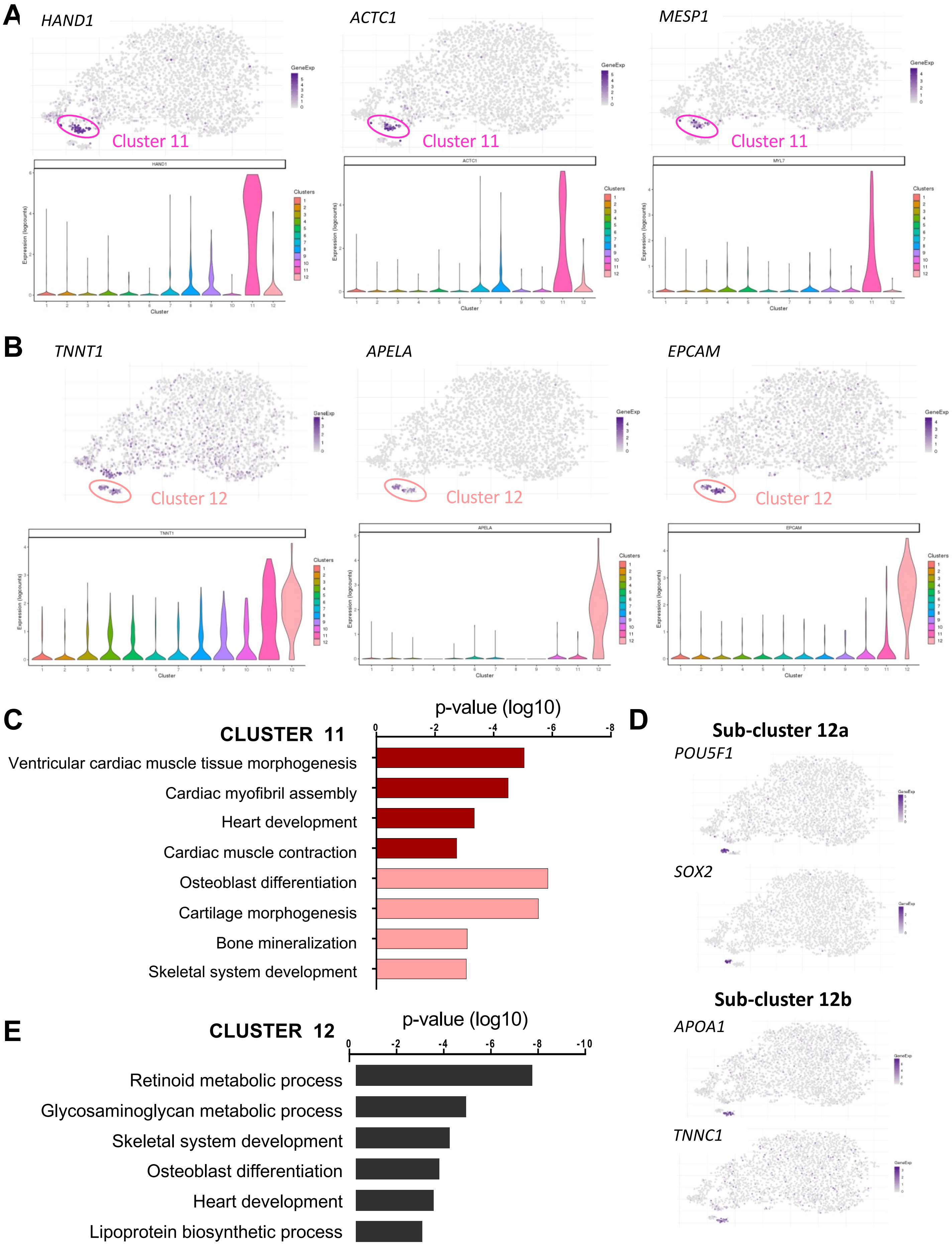
A
HAND1
ACTC1
MESP1
Cluster 11
Cluster 11
Cluster 11
B
TNNT1
APELA
EPCAM
Cluster 12
Cluster 12
Cluster 12
C
CLUSTER 11
p-value (log10)
Ventricular cardiac muscle tissue morphogenesis
Cardiac myofibril assembly
Heart development
Cardiac muscle contraction
Osteoblast differentiation
Cartilage morphogenesis
Bone mineralization
Skeletal system development
D
Sub-cluster 12a
POU5F1
SOX2
Sub-cluster 12b
APOA1
TNNC1
E
CLUSTER 12
p-value (log10)
Retinoid metabolic process
Glycosaminoglycan metabolic process
Skeletal system development
Osteoblast differentiation
Heart development
Lipoprotein biosynthetic process

**Figure S4**

**Figure S4. Characterisation of cluster 11 and 12.**

(A) t-SNE and violin plots showing the expression of *HAND1*, *ACTC1* and *MESP1* in all clusters. (B) t-SNE and violin plots showing the expression of *TNNT1*, *APELA* and *EPCAM* expression in all clusters. (C) GO terms enriched in cluster 11 upregulated genes. (D) t-SNE plots showing the expression of *POUF1*, SOX2, *APOA1* and *TNNC1* expression in all clusters. (E) GO terms enriched in cluster 12 upregulated genes.

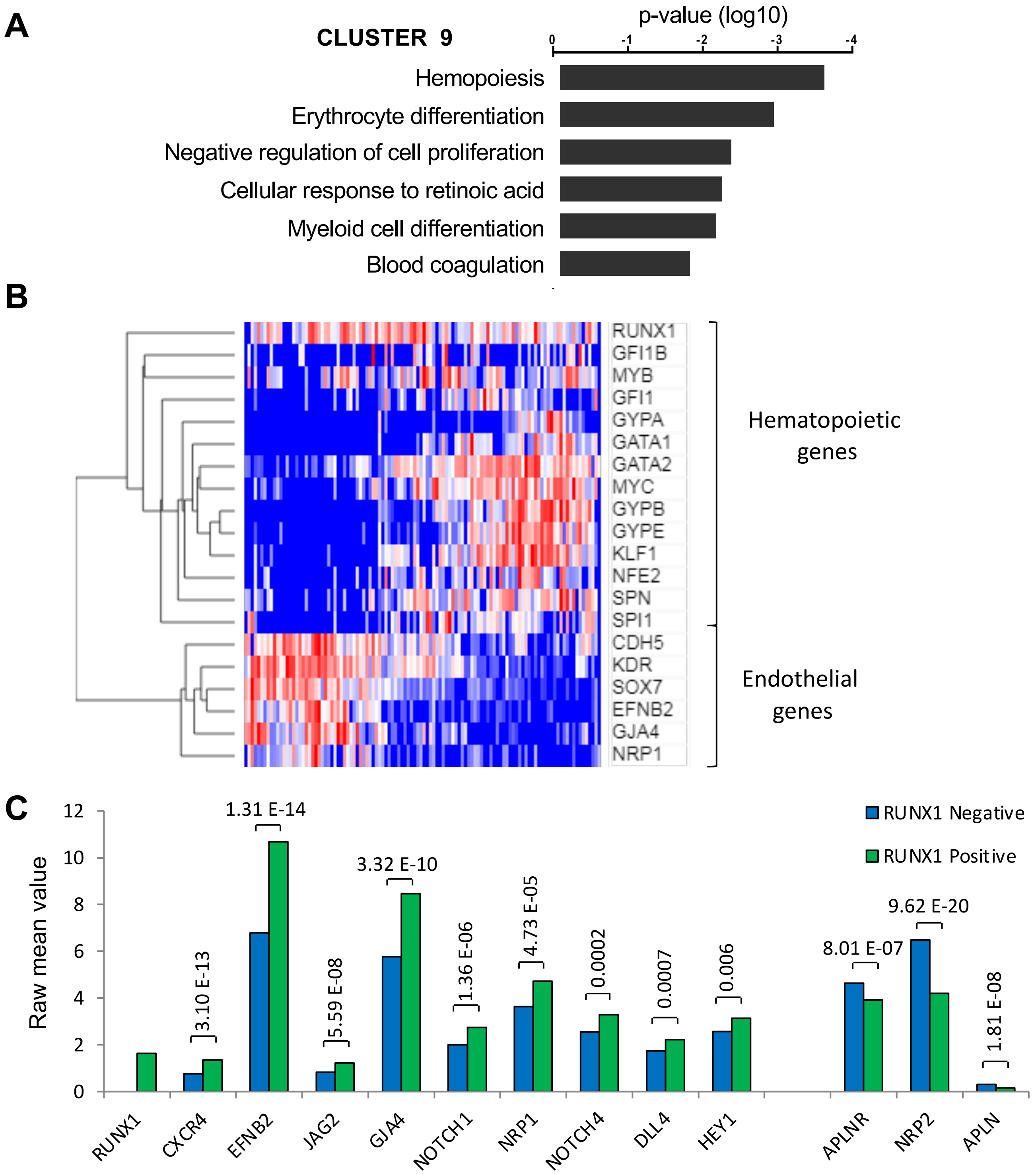
A
CLUSTER 9
p-value (log10)
0
-1
-2
-3
-4
Hemopoiesis
Erythrocyte differentiation
Negative regulation of cell proliferation
Cellular response to retinoic acid
Myeloid cell differentiation
Blood coagulation
B
RUNX1
GFI1B
MYB
GFI1
GYPA
GATA1
GATA2
MYC
GYPB
GYPE
KLF1
NFE2
SPN
SPI1
CDH5
KDR
SOX7
EFNB2
GJA4
NRP1
Hematopoietic genes
Endothelial genes
C
Raw mean value
12
10
8
6
4
2
0
RUNX1 Negative
RUNX1 Positive
3.10 E-13
1.31 E-14
5.59 E-08
3.32 E-10
1.36 E-06
4.73 E-05
0.0002
0.0007
0.006
8.01 E-07
9.62 E-20
1.81 E-08
RUNX1
CXCR4
EFNB2
JAG2
GJA4
NOTCH1
NRP1
NOTCH4
DLL4
HEY1
APLNR
NRP2
APLN

**Figure S5**

**Figure S5: Characterisation of cluster 9.**

(A) GO terms enriched in Cluster 9 upregulated genes . (B) Heatmap of unsupervised hierarchical clustering of selected endothelial and hematopoietic genes expressed in cells of cluster 9. (C) Average expression levels from scRNA-seq dataset in RUNX1-positive (green bars) and RUNX1-negative (blue bars) cells within all endothelial clusters for selected arterial and venous genes. *P*-value are indicated on the graph.

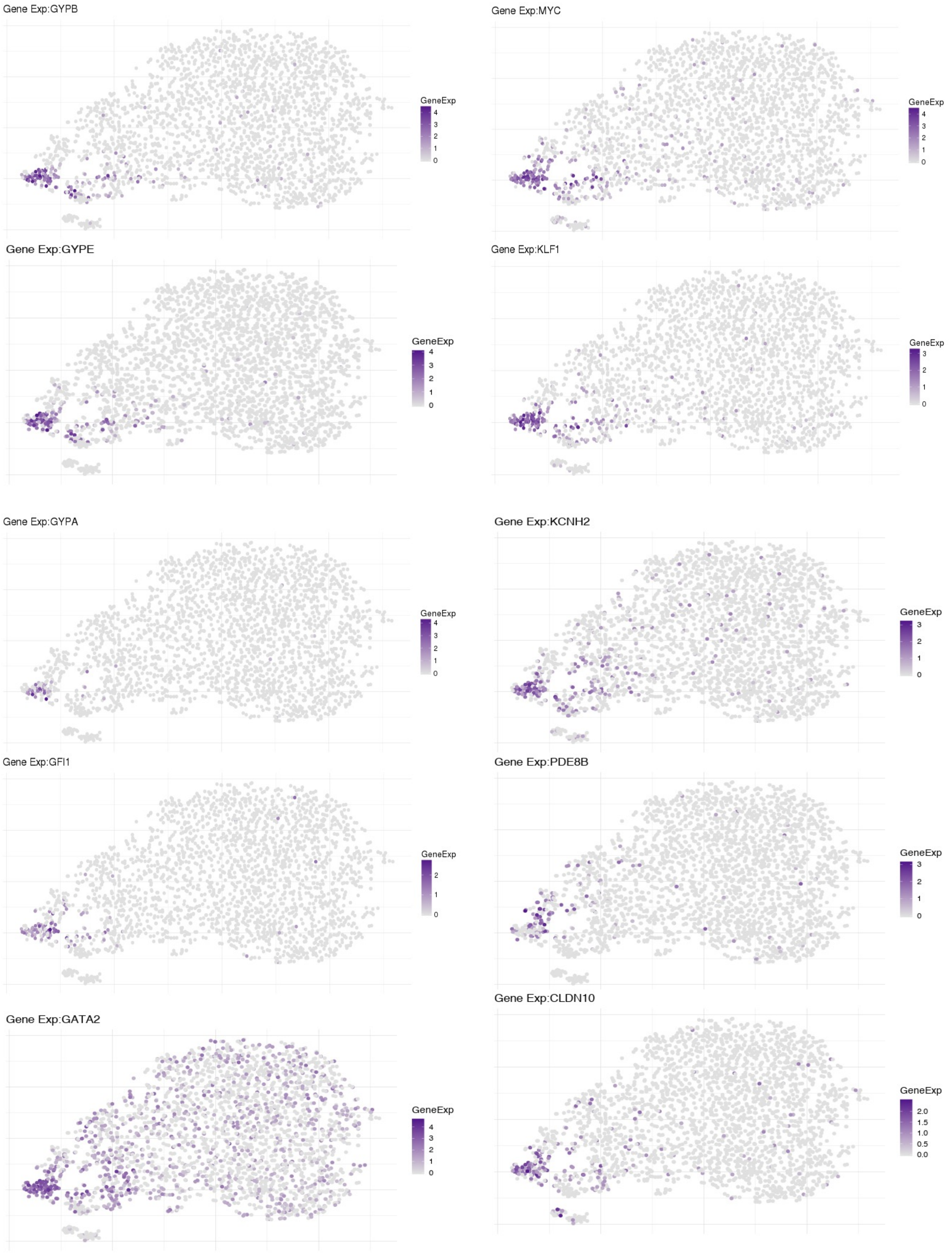

Gene Exp:GYPB
GeneExp
Gene Exp:MYC
GeneExp
Gene Exp:GYPE
GeneExp
Gene Exp:KLF1
GeneExp
Gene Exp:GYPA
GeneExp
Gene Exp:KCNH2
GeneExp
Gene Exp:GFI1
GeneExp
Gene Exp:PDE8B
GeneExp
Gene Exp:CLDN10
GeneExp
Gene Exp:GATA2
GeneExp

**Figure S6: Gene expression across clusters.**

t-SNE plots showing the expression of genes which expression is significantly upregulated in cluster 9 compared to all other clusters.

**Supplemental table 1**: Genes most  down-regulated in cluster 9 relative to all other clusters.

| Gene name | LogFC | FDR |
|---|---|---|
| SOX17 | -3.412778198 | 5.78E-47 |
| RAMP2 | -3.215549888 | 3.20E-97 |
| ADGRL4 | -2.743520292 | 6.77E-59 |
| ARHGAP29 | -2.73161832 | 5.32E-87 |
| ID1 | -2.664547243 | 1.21E-50 |
| ID3 | -2.574305153 | 1.01E-80 |
| WWTR1 | -2.538810345 | 5.96E-45 |
| LDB2 | -2.229648743 | 1.12E-48 |
| HOPX | -2.139606926 | 2.23E-46 |
| SMAGP | -2.122669477 | 4.58E-51 |
| GSN | -2.093942051 | 2.14E-47 |
| IGFBP4 | -2.077466628 | 2.69E-82 |
| IFI16 | -2.05363072 | 4.51E-66 |
| THY1 | -2.048444396 | 1.40E-58 |
| PLVAP | -1.760119338 | 1.04E-54 |
| AP1S2 | -1.69268576 | 5.76E-69 |
| VIM | -1.678153348 | 1.09E-62 |
| MARCKS | -1.603020522 | 1.68E-64 |
| TMSB10 | -1.522645529 | 1.24E-110 |
| FSCN1 | -1.452906133 | 1.98E-59 |
| RDX | -1.43984293 | 5.64E-49 |
| KDR | -1.385729208 | 3.82E-42 |
| TMSB4X | -1.382804888 | 8.86E-56 |
| SPTBN1 | -1.349716996 | 1.02E-42 |
| CALM1 | -1.178234248 | 2.29E-46 |
| MYL6 | -0.922635948 | 1.05E-42 |
